# A widely-used pollutant causes reversal of conspecific mate preference in a freshwater fish

**DOI:** 10.1101/2022.09.07.507014

**Authors:** Daniel L. Powell, Aaron D. Rose, Gil G. Rosenthal

## Abstract

Chemical communication is an important mechanism of mate choice across the animal tree of life. However, anthropogenic perturbation of the signaling environment can disrupt chemical communication and result in a breakdown of behavioral reproductive isolation. Here we find that calcium hydroxide (Ca(OH)_2_), a common and deliberately introduced anthropogenic pollutant, profoundly disrupts chemical communication in the swordtail fish *Xiphophorus birchmanni*. Moreover, it acts in a way that should promote hybridization, causing female *X. birchmanni* to prefer the chemical cues of the parapatric sister species *X. malinche*. We find that this flip in the direction of preference is attributable both to a reduced preference for conspecific signals and to a coupled strengthening of preference for sister-species signals.

## BACKGROUND

Behavioral reproductive isolation based on chemical communication is pervasive across sexually reproducing species [1–4] and can be extraordinarily fine-tuned compared to other modes of communication as a result of the diversity and specificity of odorant receptors [5,6]. In cases where individual interactions are mediated by chemical signaling, environmental disturbances that either change chemical signals or impact their receptors have the potential to disrupt olfactory signaling and break down pre-mating reproductive barriers between species [3,7,8].

Hybridization between two northern swordtail fish, *Xiphophorus birchmanni* and *X. malinche*, is thought to be mediated by disruption of chemical communication through pollutants linked to anthropogenic disturbance [9–12]. Specifically, environmentally relevant concentrations of humic acid, a common byproduct of agricultural run-off and untreated sewage, is sufficient to ablate preferences for conspecific male pheromones in female *X. birchmanni* [9]. However, many other compounds that find their way into stream water are known to affect chemical communication in fishes (reviewed in: [13,14]). Studies have largely focused on heavy metals and pesticides known to be endocrine disruptors (e.g. [7,15–19]), but other chemical compounds that do not cause mortality could also impact chemical communication. One such compound that is often deliberately introduced into aquatic environments is calcium hydroxide (Ca(OH)_2_). It is used for waste water treatment in reduction of pathogen concentrations [20–22], removal of particulate matter, and precipitation of phosphate and heavy metals [23,24].

Similarly, it is used to help combat eutrophication and acidification in lakes and aquaculture facilities [25–30]. Despite its widespread use, the effects of the addition of Ca(OH)_2_ on vertebrates remain understudied [31–33].

Over the course of more than 25 years of research in habitats of *X. birchmanni, X. malinche*, and their hybrids [10], researchers have frequently observed instances of the aftermath of calcium hydroxide being deliberately introduced into streams (Rosenthal and Powell, personal observations; Fig. S1a,b). Large quantities of powdered calcium hydroxide are introduced into riffles up-stream of pools used by humans and cattle as a method to reduce bacterial load via pH shock and flocculation [20,22,34,35]. This also creates a hostile environment for other stream organisms. In fact, because it often kills fishes at high enough doses, we have observed it used as a fishing technique (Fig. S1b,c) — similar fishing techniques are used elsewhere in México [36]. Though aquatic vertebrate mortality is high in such cases, some fish are exposed to less concentrated plumes. Field observations indicate that exposure to sublethal concentrations of calcium hydroxide may affect some fishes’ ability to perceive olfactory food cues (Supplemental information 1).

The addition of calcium hydroxide to stream water has two primary effects on water chemistry [25]. First, it increases Ca^2+^ ion concentration, which may alter the efficacy of neuronal signaling as calcium plays an important role in ORN function [37]. Second, it increases pH due to the influx of OH^-^ ions. Environmental changes in pH are known to affect chemical communication in teleost fishes [38,39]. Here, we test the effects of short-term exposure to calcium hydroxide (Ca(OH)_2_) on mate preference in the olfactory modality in female *X. birchmanni*. To distinguish the effects of Ca^2+^ ion concentrations from those of pH, we then tested the effects of calcium chloride (CaCl_2_) which raises Ca^2+^ ion concentrations but does not affect pH, and of sodium hydroxide (NaOH) which raises pH but does not affect Ca^2+^ ion concentrations.

## METHODS

### Fish collection and housing

All *X. birchmanni* (focal females and males for chemical cue production) were collected between March 2018 and January 2019 from the Río Coacuilco near the town of Coacuilco (21°5’50.85 N, 98°35’19.46 W). *X. malinche* males for chemical cue production were collected from the Chicayotla location on Río Xontla (20°55’27.24”N 98°34’34.50W; Fig. S2) [10]. All fish were housed at Texas A&M University in single sex 120 L tanks maintained at 22° C, on a 12/12 hr light cycle, and fed twice daily (Texas A&M IACUC protocol #2016-0190).

### Simultaneous olfactory signal presentation preference trials

For all tests of the female olfactory modality, mate preference was tested following a well-established protocol in which stimuli are presented simultaneously at either end of a trial lane [40,41]. Briefly, focal female *X. birchmanni* were acclimatized to a 75×19×20 cm trial lane for ten minutes after which stimulus flow at either end was initiated. Stimuli were presented for 600s. Association time, an important indicator of preference [42], with each stimulus was scored for 300s beginning once the focal female visited both association zones. To control for side-bias each female was tested a second time with the cue presentation reversed. Association times were summed across both trials for each female. Females that failed to visit both zones within 300 s in both trials were excluded from analysis (supplemental information 2). Pooled male chemical stimuli for both conspecific (*X. birchmanni*) and heterospecific (*X. malinche*) were prepared following Fisher et al. 2006 no more than 24 hrs in advance of testing (supplemental information 3) [9]. New pooled cues were prepared for each day of tests.

For assays testing the effects of calcium hydroxide on odor preference between conspecific and heterospecific stimuli, females were randomly assigned to the control (*N* = 14) or exposure group (*N* = 14) and tested on four separate days. On day zero both groups were given a sham exposure in which they were held in 4 L tanks (two per container) containing carbon filtered tap water for ten minutes prior to being placed into the trial lanes for acclimation. On day two the control group was given the same sham exposure while the exposure group was placed into identical 4 L tanks containing carbon filtered tap water and 24 mg/L calcium hydroxide for ten minutes prior to testing, a duration similar to exposures in the wild (supplemental information 4). On day four, two days after exposure, and again on day 12, ten days post-exposure, both groups were once again treated with a sham exposure containing only carbon filtered water prior to testing. A 24 mg/L concentration is similar to recommended concentrations for eutrophication remediation and 10x less than the LC50 for a closely related species, *Gambusia affinis* [25,43].

A similar set of trials using new females was performed for fish exposed to an equimolar concentration of calcium chloride (36 mg/L CaCl_2_; control *N* = 16, exposure *N* = 16; supplemental information 5). A further set of trials was run using a sodium hydroxide exposure (control *N* = 16, exposure *N* = 16) titrated to a pH of 9.21 (15.4 mg/l NaOH), equal to the calcium hydroxide exposure (supplemental information 5). For both of these trial sets, day 12 was omitted as females had recovered species typical responses by day four.

### Olfactory signal versus water control preference trials

Calcium hydroxide exposure may have a singular effect on *X. birchmanni* female responses to either conspecific or heterospecific male cues, or it may affect responses to both. To address this question, we tested a new set of females (control *N* = 16, exposure *N* = 16) for conspecific (*X. birchmanni*) chemical cue against a water blank, and heterospecific (*X. malinche*) cues against the same. Each female was tested twice for each combination to control for side bias (four tests total per fish). The order and side of cue presentation were randomized. Trials were performed as described above for Ca(OH)_2_ exposure.

### Statistical analysis

To compare chemical trials within treatments (control, and exposure) across all time points we used separate one way repeated measures ANOVAs on net preference for each chemical exposure (Ca(OH)_2_, CaCl_2_, and NaOH). Where residual distributions or variances did not meet the assumption of normality or equality a Wilcoxon signed-rank tests or Kruskal-Wallis rank-sum tests were substituted. All analyses were done in the R computing environment.

To assess female preference in all chemical trials within a given group (control or exposure) for a given timepoint we used two-tailed, one sample t-tests for mean net association time with the null expectation of no preference.

## RESULTS

Responsiveness between control and exposure fish did not differ for any trial (Table S1). Tests comparing net association times for any given day for all trials are summarized in table S2.

### Simultaneous olfactory signal presentation preference trials

Response rates did not differ between control and exposure groups for any test (Table S1; Fisher’s exact test, *p* > 0.1). Net association time with the conspecific olfactory cue did not differ significantly for control females across the Ca(OH)_2_ trials (Fig. 1a; repeated measures ANOVA, *F*(3, 39) = 0.226, *p* = 0.878). However, net association time for the Ca(OH)_2_ exposed fish did differ significantly across time points (Fig 1b; repeated measures ANOVA *F*(3,45) = 7.875, *p* = 0.00025) with net association time with the conspecific cue being lower on the day of and two days after exposure (days two and four) than two days prior to (day 0) and ten days after (day 12) exposure (Fig. 1b; TukeyHSD, *p* < 0.05). Unexpectedly, on the day of exposure (day two; *N* _responsive_ = 11, two tailed, one sample *t-*test, *t* = -2.27 *p* = 0.046) and two days after exposure (day 4; *N* _responsive_ = 12, two tailed, one sample *t-*test, *t* = -2.77, *p* = 0.018) the mean net association times were significantly negative representing a switch in preference to heterospecific male chemical cues (Fig. 1b). By 10 days post exposure (day 12) the species typical conspecific preference was recovered (*N* _responsive_ = 14, two tailed Wilcoxon signed-rank test, *p* = 0.0067).

**Figure 1.**
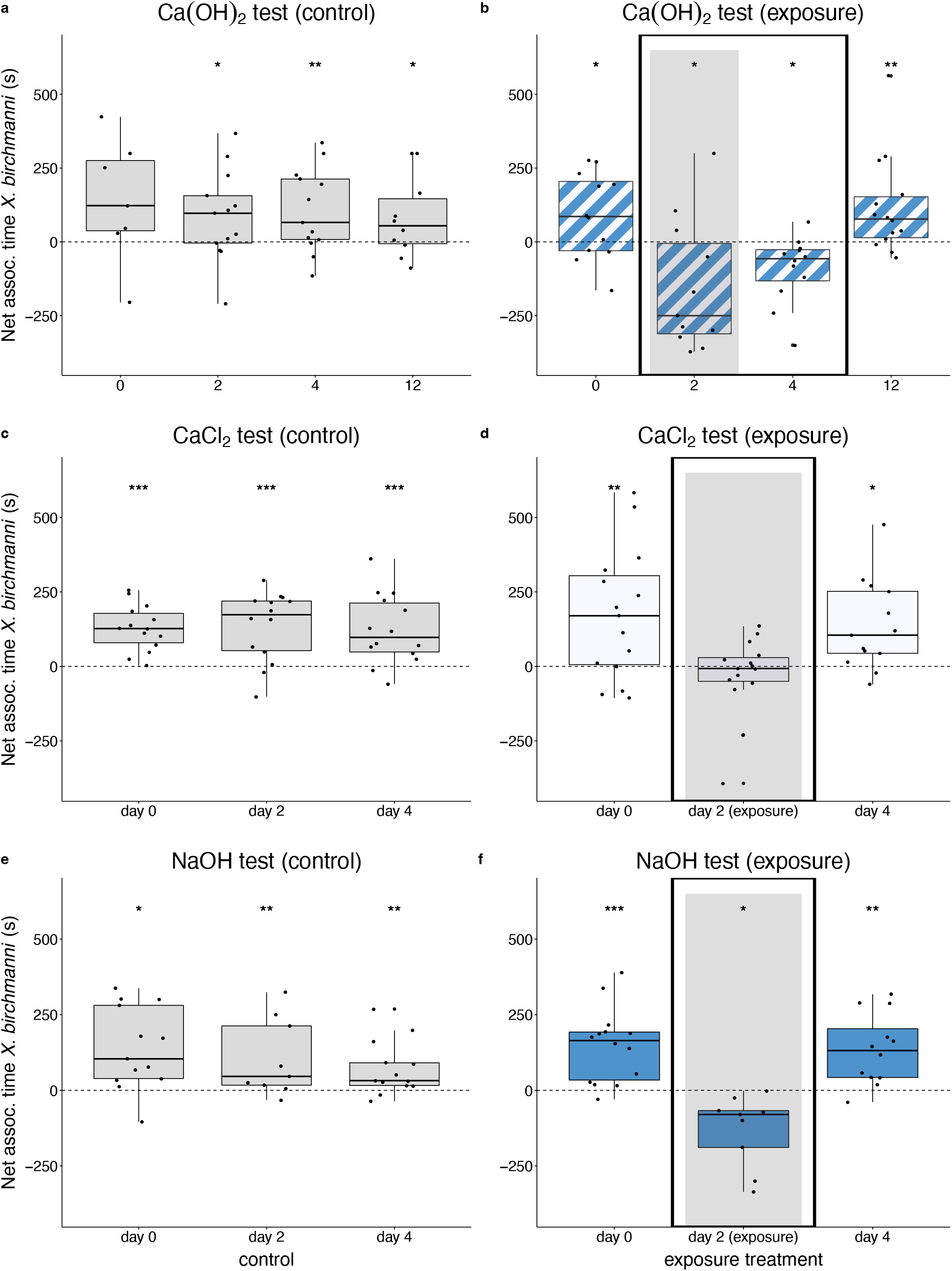
Effect of chemical exposure on female *X. birchmanni* preference for male chemical stimuli. Net association time of female *X. birchmanni* with conspecific (*X. birchmanni*) over heterospecific (*X. malinche*) male chemical stimuli for Ca(OH)_2_ exposure trials, a) control treatment and b) exposure treatment; CaCl_2_ exposure trials, c) control treatment and d) exposure treatment; and NaOH exposure trials, e) control treatment and, f) exposure treatment. Shaded columns represent trials initiated immediately after experimental chemical exposure. Bold black boxes surround trials with significantly different female responses relative to other trials (trials not included in boxed area; TukeyHSD, *p* < 0.05). Asterisks represent mean association times different from zero (one-sample, * *p* < 0.05, ** *p* < 0.01, *** *p* < 0.001).

Net association time with the conspecific olfactory cue did not differ significantly for control females across the CaCl_2_ trials (Fig. 1c; repeated measures ANOVA, *F*(2,39) = 0.059, *p* = 0.942). However, net association time for the CaCl_2_ exposed fish did differ significantly across time points with net association time with the conspecific cue being less on the day of exposure (day 2) than two days prior to (day 0) and two days post (day 4) exposure (Fig. 1d; repeated measures ANOVA, *F*(2,40) = 5.854, *p* = 0.006; TukeyHSD, *p* < 0.038). In this case, on the day of exposure there was no significant preference (Fig. 1d; *N* _responsive_ = 15, Wilcoxon signed-rank test, *p* = 0.720)

As with the previous two sets of trials, net association time with the conspecific olfactory cue did not differ significantly for control females across the NaOH trials (Fig 1e; Kruskal-Wallis rank sum, chi-squared = 2.44, *p* = 0.296, d.f. = 2). Responses of NaOH exposed females were qualitatively similar to the Ca(OH)_2_ exposed fish with net conspecific preference being significantly less on the day of exposure (day 2) than two day prior to (day 0) or two days post (day 4) exposure (Fig. 1f; repeated measures ANOVA, *F*(2, 32) = 17.29, *p* = 0.0000081; TukeyHSD, *p* < 0.00055). As with Ca(OH)_2_ exposed females, on the day of exposure (day 2) females subjected to NaOH significantly preferred the heterospecific cue (*N* _responsive_ = 9, two tailed one sample *t-*test, *t* = -3.283, *p* = 0.0111). Species typical conspecific cue preference was recovered by day 4 (Fig. 1f; *N* _responsive_ = 12, two tailed one sample *t-*test, *t* = 3.993, *p* = 0.0021).

### Olfactory signal versus water control preference trials

Control females showed significant mean net association time for conspecific chemical cues over a water blank (Fig. 2a; *N* _responsive_ = 14, two tailed one sample *t-*test, *t* = 3.233, *p* = 0.0065), but no preference for heterospecific cues over water (Fig. 2a; *N* _responsive_ = 13, two tailed one sample *t-* test, *t* = -1.938, *p* = 0.187). Ca(OH)_2_ exposed females showed no preference for conspecific cues over water (Fig. 2b; *N* _responsive_ = 12, two tailed one sample *t-*test, *t* = -0.994, *p* = 0.342) but a strong preference for heterospecific cues over water (Fig 2b; *N* _responsive_ = 16, two tailed one sample *t-*test, *t* = 2.979, *p* = 0.0094).

**Figure 2.**
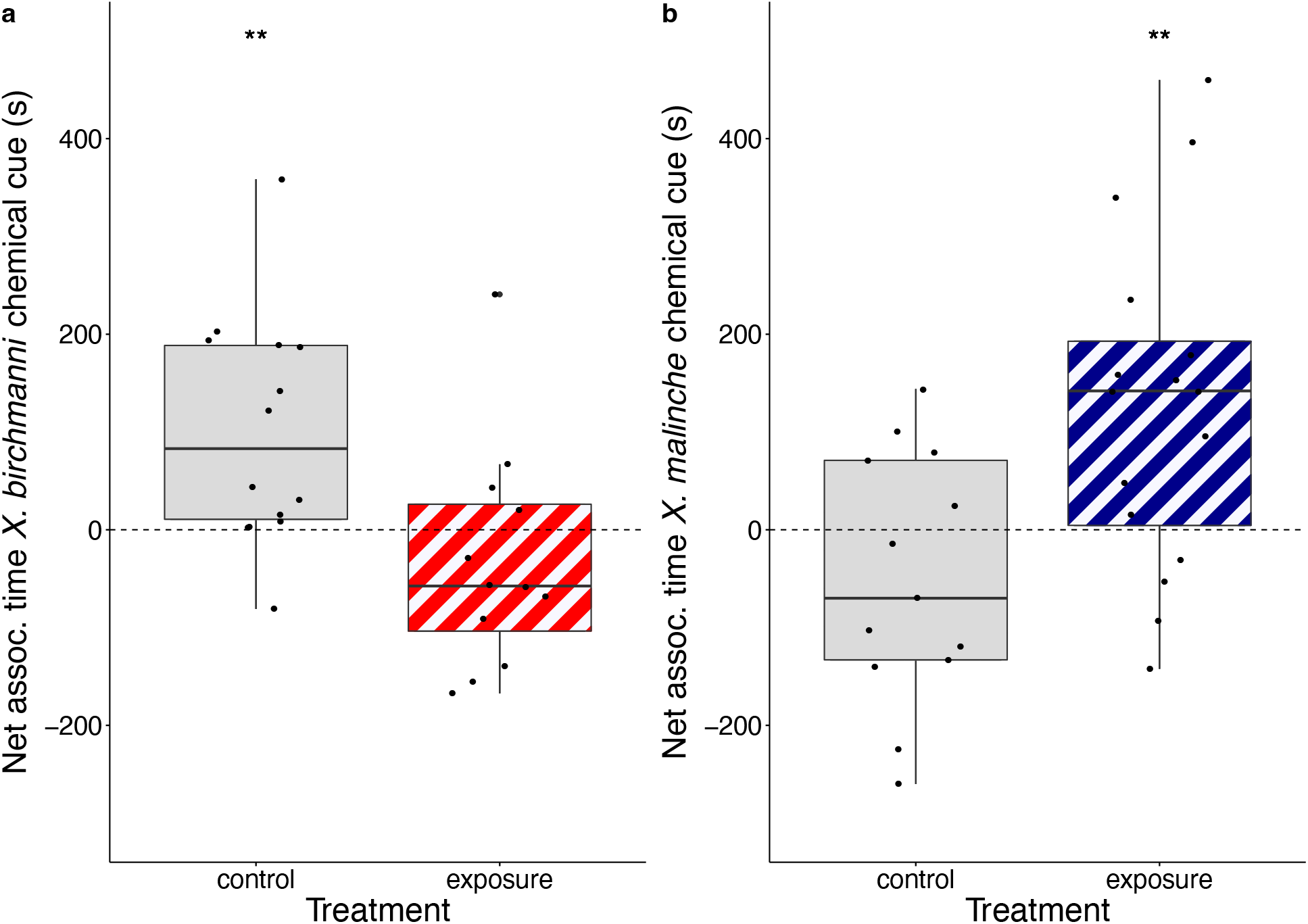
Female preference for olfactory signals compared to water blanks. Net association time of control and Ca(OH)_2_ exposed female *X. birchmanni* for a) conspecific (*X. birchmanni*; red diagonal box) male chemical stimuli over a water blank (grey box) and b) heterospecific (*X. malinche*; blue diagonal box) male chemical stimuli over a water blank (grey box). Boxes represent interquartile range, horizontal bar within boxes represents median, whiskers represent minima and maxima excluding outliers, and dots represent raw data. Asterisks represent mean association times different from zero (one sample, two-tailed t-test, ** *p* < 0.01).

## DISCUSSION

In this study, female mating preference assays show that not only can a common, deliberately introduced anthropogenic pollutant disrupt chemical communication in *X. birchmanni*, but it does so in a way that should promote hybridization. When exposed to low concentrations of calcium hydroxide, strong conspecific preference typically shown by *X. birchmanni* females is not just abolished, but rather a preference for the sympatric sister species *X. malinche* emerges, an effect which lasts for at least two days (Fig. 1b). This unexpected switch in to heterospecific preference could lead directly to interspecific matings following a disturbance [44]. Calcium ions alone were not sufficient to cause the switch in preference seen in the calcium hydroxide-exposed fish. However, an increased Ca^++^ concentration seems to interact with the olfactory periphery given that exposure eliminated preference in the olfactory modality while not affecting motivation to mate, as exposure females were as responsive to male cues as controls (Table S1; Fisher’s exact test, *p* = 1). Raising the pH using NaOH had a qualitatively similar effect to calcium hydroxide exposure, suggesting that other compounds that affect pH may have important effects on chemical signaling (Fig. 1f).

Calcium hydroxide-exposed *X. birchmanni* females showed no preference for conspecific cues when tested against a water blank but strongly preferred *X. malinche* (Fig. 2), whereas control fish preferred conspecific cues over water and were indifferent toward heterospecific cues (Fig. 2a). These results indicate that the preference reversal after exposure in the simultaneous choice trials is explained by the interaction of flips in hedonic value assigned to both species-specific cues (i.e. from positive to neutral for the *X. birchmanni* cue and neutral to positive for the *X. malinche* cue). Chemoreceptors in *Xiphophorus* have not been characterized, but given the strong olfactory based species recognition exhibited by *X. birchmanni* under natural conditions [9,45–48] it seems reasonable that the preference changes shown here are related to the interaction of Ca(OH)_2_ exposed chemoreceptors and components of species specific pheromone profiles [49]. Given that we used the full gamut of odorant cues left behind by multiple males interacting with females and with each other, it is also possible that the treated fish are more responsive to other cues present in the stimulus water that are not associated with mate preference.

There are three likely pathways by which altering the chemical environment can affect olfactory communication. First, it can change the composition or conformation of the signal molecules [50]. Second, it can affect the response sensitivity of olfactory receptor neurons (ORNs) to signal molecules [38,51–53]. Third, it may alter signal processing down-stream of the sensory periphery [2]. This study does not directly exclude the first scenario, but due to the intermittent pulsed nature of this point source pollution and the flip in hedonic value still evident two days post exposure, increased calcium hydroxide concentration is likely to affect signal perception or down-stream neural processing rather than signal molecules themselves.

Disentangling the second and third scenarios will be an interesting avenue for future exploration.

Though not explicitly tested, the ten-day post exposure recovery is faster than reported ORN regeneration in other vertebrates including teleost fishes, suggesting that exposure at these concentrations does not cause neuronal degeneration [54–56]. That strong preferences were still exhibited after exposure to two of the three chemicals tested, further suggests that the ORNs were not ablated.

## CONCLUSIONS

Hybridization ultimately requires the breakdown of reproductive barriers, an important first step of which can be individuals choosing heterospecific mates. Here we show that an inorganic compound generally thought to be safe for aquatic environments and used actively in remediation can not only disrupt chemical communication in these closely related sympatric swordtails, but actually causes a switch in preference from the conspecific to heterospecific, a scenario that may promote ongoing gene flow. This study adds to the growing body of evidence that sublethal concentrations of pollutants can have unforeseen ecological and evolutionary consequences [8,9,57–60].

## Acknowledgements

We thank members of the Rosenthal and Schumer labs for helpful discussion and/or feedback on earlier versions of this work and Jeffery White for help running behavioral trials. We are grateful to the Mexican federal government for permission to collect samples (Permiso de Pesca de Fomento no. PPF/DGOPA 173/14 and Permiso de Pesca de Fomento no. PPF/DGOPA-002/19). This work was supported by NSF GRFP 2019273798 to D.L. Powell and NSF IOS-1354172 and NSF IOS-1755327 to G.G. Rosenthal.

## Author contributions

D.L. Powell, and G.G. Rosenthal conceived of this project. D.L. Powell, and A. Rose collected and analyzed data. All authors wrote the manuscript.

## Data availability

All raw behavior data will be deposited in DryadXXXXX.

## REFERENCES

1. Smadja C, Butlin RK. 2008 On the scent of speciation: the chemosensory system and its role in premating isolation. Heredity 102, 77. (doi:10.1038/hdy.2008.55)

2. Rosenthal GG. 2018 Evaluation and hedonic value in mate choice. Current Zoology 64, 485–492. (doi:10.1093/cz/zoy054)

3. Wyatt TD. 2014 Pheromones and animal behavior: chemical signals and signatures. 2nd edn. Cambridge University Press.

4. Brock CD, Wagner CE. 2018 The smelly path to sympatric speciation? Molecular Ecology 27, 4153–4156. (doi:10.1111/mec.14845)

5. Xue B, Rooney AP, Kajikawa M, Okada N, Roelofs WL. 2007 Novel sex pheromone desaturases in the genomes of corn borers generated through gene duplication and retroposon fusion. Proceedings of the National Academy of Sciences 104, 4467–4472. (doi:10.1073/pnas.0700422104)

6. Leary GP, Allen JE, Bunger PL, Luginbill JB, Linn CE, Macallister IE, Kavanaugh MP, Wanner KW. 2012 Single mutation to a sex pheromone receptor provides adaptive specificity between closely related moth species. Proceedings of the National Academy of Sciences 109, 14081–14086. (doi:10.1073/pnas.1204661109)

7. van der Sluijs I et al. 2011 Communication in troubled waters: responses of fish communication systems to changing environments. Evolutionary Ecology 25, 623–640. (doi:10.1007/s10682-010-9450-x)

8. Candolin U. 2019 Mate choice in a changing world. Biological Reviews 94, 1246–1260. (doi:10.1111/brv.12501)

9. Fisher HS, Wong BBM, Rosenthal GG. 2006 Alteration of the chemical environment disrupts communication in a freshwater fish. Proceedings of the Royal Society B: Biological Sciences 273, 1187–1193. (doi:10.1098/rspb.2005.3406)

10. Rosenthal GG, de la Rosa Reyna XF, Kazianis S, Stephens MJ, Morizot DC, Ryan MJ, García de León FJ. 2003 Dissolution of sexual signal complexes in a hybrid zone between the swordtails Xiphophorus birchmanni and Xiphophorus malinche (Poeciliidae). Copeia 2003, 299–307. (doi:10.1643/0045-8511(2003)003[0299:Dossci]2.0.Co;2)

11. Hankison SJ, Morris MR. 2003 Avoiding a compromise between sexual selection and species recognition: female swordtail fish assess multiple species-specific cues. Behavioral Ecology 14, 282–287. (doi:10.1093/beheco/14.2.282)

12. Crapon de Caprona MD, Ryan MJ. 1990 Conspecific mate recognition in swordtails, Xiphophorus nigrensis and X. pygmaeus (Poeciliidae): olfactory and visual cues. Animal Behaviour 39, 290–296. (doi:10.1016/S0003-3472(05)80873-5)

13. Lürling M, Scheffer M. 2007 Info-disruption: pollution and the transfer of chemical information between organisms. Trends in Ecology & Evolution 22, 374–379. (doi:10.1016/j.tree.2007.04.002)

14. Klaprat DA, Evans RE, Hara TJ. 1992 Environmental contaminants and chemoreception in fishes. In Fish Chemoreception (ed TJ Hara), pp. 321–341. Dordrecht: Springer Netherlands. (doi:10.1007/978-94-011-2332-7_15)

15. Tomkins P, Saaristo M, Allinson M, Wong BBM. 2016 Exposure to an agricultural contaminant, 17β-trenbolone, impairs female mate choice in a freshwater fish. Aquatic Toxicology 170, 365–370. (doi:10.1016/j.aquatox.2015.09.019)

16. Honda RT, Fernandes-de-Castilho M, Val AL. 2008 Cadmium-induced disruption of environmental exploration and chemical communication in matrinxã, Brycon amazonicus. Aquatic Toxicology 89, 204–206. (doi:10.1016/j.aquatox.2008.07.001)

17. Lee Y-M, Seo JS, Kim I-C, Yoon Y-D, Lee J-S. 2006 Endocrine disrupting chemicals (bisphenol A, 4-nonylphenol, 4-tert-octylphenol) modulate expression of two distinct cytochrome P450 aromatase genes differently in gender types of the hermaphroditic fish Rivulus marmoratus. Biochemical biophysical research communications 345, 894–903. (doi:doi.org/10.1016/j.bbrc.2006.04.137)

18. Ward AJ, Duff AJ, Horsfall JS, Currie S. 2007 Scents and scents-ability: pollution disrupts chemical social recognition and shoaling in fish. Proceedings of the Royal Society B: Biological Sciences 275, 101–105. (doi:doi.org/10.1098/rspb.2007.1283)

19. Sárria M, Santos M, Reis-Henriques M, Vieira N, Monteiro N. 2011 The unpredictable effects of mixtures of androgenic and estrogenic chemicals on fish early life. Environment international 37, 418–424. (doi:doi.org/10.1016/j.envint.2010.11.004)

20. Grabow WO, Middendorff IG, Basson NC. 1978 Role of lime treatment in the removal of bacteria, enteric viruses, and coliphages in a wastewater reclamation plant. Appl Environ Microbiol 35, 663–669. (doi:10.1128/aem.35.4.663-669.1978)

21. Gerba CP. 1981 Virus Survival in Wastewater Treatment. In Viruses and Wastewater Treatment (eds M Goddard, M Butler), pp. 39–48. Pergamon. (doi:10.1016/B978-0-08-026401-1.50011-7)

22. Reinthaler FF et al. 2010 ESBL-producing E. coli in Austrian sewage sludge. Water Research 44, 1981–1985. (doi:10.1016/j.watres.2009.11.052)

23. del Bubba M, Checchini L, Pifferi C, Zanieri L, Lepri L. 2004 Olive Mill Wastewater Treatment by a Pilot-Scale Subsurface Horizontal Flow (SSF-h) Constructed Wetland. Annali di Chimica 94, 875–887. (doi:10.1002/adic.200490110)

24. Semerjian L, Ayoub GM. 2003 High-pH–magnesium coagulation–flocculation in wastewater treatment. Advances in Environmental Research 7, 389–403. (doi:10.1016/S1093-0191(02)00009-6)

25. Murphy T, Hall K, Northcote T. 1988 Lime treatment of a hardwater lake to reduce eutrophication. Lake Reservoir Management 4, 51–62. (doi:10.1080/07438148809354813)

26. Murphy TP, Prepas EE, Lim JT, Crosby JM, Walty DT. 1990 Evaluation of Calcium Carbonate and Calcium Hydroxide Treatments of Prairie Drinking Water Dugouts. Lake and Reservoir Management 6, 101–108. (doi:10.1080/07438149009354700)

27. Fraser JE, Britt DL. 1982 Liming of acidified waters: a review of methods and effects on aquatic ecosystems. Final report., 208.

28. Johnson DB, Hallberg KB. 2005 Acid mine drainage remediation options: a review. Science of The Total Environment 338, 3–14. (doi:10.1016/j.scitotenv.2004.09.002)

29. Clair TA, Hindar A. 2005 Liming for the mitigation of acid rain effects in freshwaters: a review of recent results. Environmental Reviews 13, 91–128. (doi:10.1139/a05-009)

30. Furtado PS, Poersch LH, Wasielesky W. 2011 Effect of calcium hydroxide, carbonate and sodium bicarbonate on water quality and zootechnical performance of shrimp Litopenaeus vannamei rearsed in bio-flocs technology (BFT) systems. Aquaculture 321, 130–135. (doi:10.1016/j.aquaculture.2011.08.034)

31. Miskimmin BM, Donahue WF, Watson D. 1995 Invertebrate community response to experimental lime (Ca(OH)2) treatment of an eutrophic pond. Aquatic Sciences 57, 20–30. (doi:10.1007/bf00878024)

32. Leoni B, Morabito G, Rogora M, Pollastro D, Mosello R, Arisci S, Forasacco E, Garibaldi L. 2007 Response of planktonic communities to calcium hydroxide addition in a hardwater eutrophic lake: results from a mesocosm experiment. Limnology 8, 121–130. (doi:10.1007/s10201-007-0202-8)

33. Ghadouani A, Alloul BP, Zhang Y, Prepas, E E. 1998 Relationships between zooplankton community structure and phytoplankton in two lime-treated eutrophic hardwater lakes. Freshwater Biology 39, 775–790. (doi:10.1046/j.1365-2427.1998.00318.x)

34. Lucena F, Duran AE, Morón A, Calderón E, Campos C, Gantzer C, Skraber S, Jofre J. 2004 Reduction of bacterial indicators and bacteriophages infecting faecal bacteria in primary and secondary wastewater treatments. Journal of applied microbiology 97, 1069–1076. (doi:10.1111/j.1365-2672.2004.02397.x)

35. Grupo Calidra Química Natural. 2000 Manual de usos ecológicos de la cal. Grupo Calidra Química natural. See https://books.google.it/books?id=ixsYogEACAAJ.

36. Tobler M, Culumber ZW, Plath M, Winemiller KO, Rosenthal GG. 2011 An indigenous religious ritual selects for resistance to a toxicant in a livebearing fish. Biology Letters 7, 229–232. (doi:10.1098/rsbl.2010.0663)

37. Nicholls JG, Martin AR, Wallace BG, Fuchs PA. 2001 From neuron to brain. Sinauer Associates Sunderland, MA.

38. Dew WA, Pyle GG. 2014 Smelling salt: Calcium as an odourant for fathead minnows. Comparative Biochemistry and Physiology Part A: Molecular & Integrative Physiology 169, 1–6. (doi:10.1016/j.cbpa.2013.12.005)

39. Heuschele J, Candolin U. 2007 An increase in pH boosts olfactory communication in sticklebacks. Biology Letters 3, 411–413. (doi:10.1098/rsbl.2007.0141)

40. McLennan DA, Ryan MJ. 1999 Interspecific recognition and discrimination based upon olfactory cues in northern swordtails. Evolution 53, 880–888. (doi:10.2307/2640728)

41. McLennan DA, Ryan MJ. 1997 Responses to conspecific and heterospecific olfactory cues in the swordtail Xiphophorus cortezi. Animal Behaviour 54, 1077–1088. (doi:10.1006/anbe.1997.0504)

42. Walling CA, Royle NJ, Lindström J, Metcalfe NB. 2010 Do female association preferences predict the likelihood of reproduction? Behavioral Ecology Sociobiology 64, 541–548. (doi:10.1007/s00265-009-0869-4)

43. Wallen I, Greer W. 1957 Toxicity to Gambusia Affinis of Certain Pure Chemicals in Turbid Waters. Sewage and Industrial Wastes 29, 695–711.

44. Rosenthal GG. 2013 Individual mating decisions and hybridization. Journal of Evolutionary Biology 26, 252–255. (doi:10.1111/jeb.12004)

45. Verzijden MN, Culumber ZW, Rosenthal GG. 2012 Opposite effects of learning cause asymmetric mate preferences in hybridizing species. Behavioral Ecology 23, 1133–1139. (doi:10.1093/beheco/ars086)

46. Fisher HS, Rosenthal GG. 2006 Hungry females show stronger mating preferences. Behavioral Ecology 17, 979–981. (doi:10.1093/beheco/arl038)

47. Fisher HS, Rosenthal GG. 2006 Female swordtail fish use chemical cues to select well-fed mates. Animal Behaviour 72, 721–725. (doi:10.1016/j.anbehav.2006.02.009)

48. Wong BBM, Fisher HS, Rosenthal GG. 2005 Species recognition by male swordtails via chemical cues. Behavioral Ecology 16, 818–822. (doi:10.1093/beheco/ari058)

49. Cui R, Delclos PJ, Schumer M, Rosenthal GG. 2017 Early social learning triggers neurogenomic expression changes in a swordtail fish. Proceedings of the Royal Society B: Biological Sciences 284, 20170701. (doi:10.1098/rspb.2017.0701)

50. Hubbard PC, Barata EN, Canario AVM. 2002 Possible disruption of pheromonal communication by humic acid in the goldfish, Carassius auratus. Aquatic Toxicology 60, 169–183. (doi:10.1016/S0166-445X(02)00002-4)

51. Burnard D, Gozlan RE, Griffiths SW. 2008 The role of pheromones in freshwater fishes. Journal of Fish Biology 73, 1–16. (doi:10.1111/j.1095-8649.2008.01872.x)

52. Lazzari M, Bettini S, Milani L, Maurizii MG, Franceschini V. 2 Differential response of olfactory sensory neuron populations to copper ion exposure in zebrafish. Aquatic Toxicology 183, 54–62. (doi:10.1016/j.aquatox.2016.12.012)

53. Reisert J, Matthews HR. 2001 Response properties of isolated mouse olfactory receptor cells. Journal of Physiology 530, 113–122. (doi:10.1111/j.1469-7793.2001.0113m.x)

54. Cancalon P. 1982 Degeneration and regeneration of olfactory cells induced by ZnSO4 and other chemicals. Tissue and Cell 14, 717–733. (doi:10.1016/0040-8166(82)90061-1)

55. Graziadei PPC, Graziadei GAM. 1979 Neurogenesis and neuron regeneration in the olfactory system of mammals. I. Morphological aspects of differentiation and structural organization of the olfactory sensory neurons. Journal of Neurocytology 8, 1–18. (doi:10.1007/bf01206454)

56. Zippel HP. 2000 In goldfish the discriminative ability for odours persists after reduction of the olfactory epithelium, and rapidly returns after olfactory nerve axotomy and crossing bulbs. J Philosophical Transactions of the Royal Society of London. Series B: Biological Sciences 355, 1219–1223. (doi:10.1098/rstb.2000.0671)

57. Dudgeon D et al. 2006 Freshwater biodiversity: importance, threats, status and conservation challenges. Biological Reviews 81, 163–182. (doi:10.1017/S1464793105006950)

58. Hayes T, Haston K, Tsui M, Hoang A, Haeffele C, Vonk A. 2002 Herbicides: feminization of male frogs in the wild. Nature 419, 895. (doi:10.1038/419895a)

59. Crispo E, Moore J, Lee-Yaw JA, Gray SM, Haller BC. 2011 Broken barriers: human-induced changes to gene flow and introgression in animals: an examination of the ways in which humans increase genetic exchange among populations and species and the consequences for biodiversity. BioEssays 33, 508–518. (doi:10.1002/bies.201000154)

60. Michelangeli M, Martin JM, Pinter-Wollman N, Ioannou CC, McCallum ES, Bertram MG, Brodin T. 2022 Predicting the impacts of chemical pollutants on animal groups. Trends in Ecology & Evolution (doi:10.1016/j.tree.2022.05.009)

