## Supplementary information for "A widely-used pollutant causes reversal of conspecific mate preference in a freshwater fish"

Daniel L. Powell, Aaron Rose, Gil G. Rosenthal

### 1. Observations of $Ca(OH)_2$ usage in *Xiphophorus* habitats

On several collecting trips between 2012 and 2019 we witnessed the aftermath of deliberate $Ca(OH)_2$ introduction in the Río Huazalingo and Calnali drainages. These events were evident by the presence of white powder lining the riffle banks at the site of introduction, littered $Ca(OH)_2$ bags near-by, discolored periphyton and dead or dying fish immediately down- stream (Fig. S1). On two separate occasions in the Río Huazalingo drainage we DLP observed surviving swordtails — pure *Xiphophorus malinche* in instance and *X. malinche* x *X. birchmanni* hybrids in the other — feeding vigorously on the $Ca(OH)_2$-affected periphyton directly downstream of several baited minnow traps. Despite being directly in the scent plume of the bait, no fish entered the traps over the course of ~1 hour. Traps deployed upstream of the $Ca(OH)_2$ introduction caught many swordtails on both occasions. Based on these observations we hypothesized that exposure to non-lethal concentrations of $Ca(OH)_2$ may affect olfaction in swordtails as the observed fish seemed motivated to feed but were not attracted by the dog food — a time tested swordtail attractant — inside the traps. Though we do not know precisely how long before our collection attempts these sites were dosed, it is likely based on rainfall patterns that the event took place no earlier than 48 hrs prior.

### 2. Simultaneous olfactory signal presentation preference trials

For all olfactory tests, mate preference was tested following a well-established protocol in which two males stimuli are presented simultaneously at either end of a trial lane [1,2].

Trials were conducted by placing a female *X. birchmanni* in a 75x19x20 cm acrylic trial lane and allowing her to acclimate for ten minutes. Lanes were divided into three sections of equal size (one association zone on either side where stimuli were presented and a central neutral zone containing a small acrylic shelter). Each lane was fitted with a stimulus delivery system at the end of each association zone run by peristaltic pumps. After the ten-minute acclimatization period, stimulus flow was initiated and delivered for 600s. Flow rate was set to 1.2 ml/min and regulated with medical-grade IV flow regulators. To control for side-bias each female was tested a second time with the cue presentation reversed. Water was changed and lanes were cleaned between trials.

Once a female visited all three zones following the initiation of stimuli flow, time spent in each stimulus association zone was recorded for 300s. Association time with a given stimulus has been shown to be a robust proxy for realized mate choices in swordtail fishes [3]. Association times were summed across both trials for each female. Females that failed to visit both association zones within 300s in both tests were scored as unresponsive and excluded from analysis.

**3. Pooled male stimulus production**

Male chemical stimuli for both conspecific (*X. birchmanni*) and heterospecifics (*X. malinche*) were prepared by placing five male fish in a 22 L tank filled with 20 liters of previously aerated and carbon filtered tap water for four hours following Fisher et al. 2006 [4]. In order to elicit release of urine-born pheromones, visual cues of females were provided by an adjacent tank containing five conspecific females. Cue production tanks were equipped with a tight fittings glass lids, an air stone for circulation, and small acrylic structures for cover. All tanks and associated equipment were cleaned with detergent and water, silicone seams were bathed with hydrogen peroxide to remove pheromone residues, and thoroughly rinsed between rounds of cue production. Cues were prepared no more than 24 hrs in advance of testing, and new cue was prepared for each day of tests.

**4. Surface waterflow measurements for exposure time estimates**

The pulsed and ephemeral nature of the calcium hydroxide pollution events described in the main text and in supplemental information 1 suggests that the physiology of the receiver (i.e. the nasal epithelia of females) is likely affected rather than the urine-borne signal itself which could be released by courting males well after the hydro-chemical effects of the $Ca(OH)_2$ have dissipated. As such we chose to expose the focal females ahead of but not during the preference trials leaving the actual male cues unaffected by our chemical manipulations. The 10-minute duration represents a rough estimation of typical breeding season flow time through pools in the Río Calnali containing hybrid populations in the wild. Mean center-of-stream surface flow rate was $0.013 \pm 0.003$ m/s for ten pools measured by float time of a tennis ball over a 10 meter distance (3 replicates per pool; 30 measurements across 10 pools total) in May 2018.

**6. pH and chemical manipulation of source water**

The pH of Río Coacuilco river water in January 2019 at the time the last focal female *X. birchmanni* were collected was 8.2. The Texas A&M University dechlorinated tap-water used for everyday husbandry as well as all behavioral trials described in this manuscript and to which all focal fish were acclimated for at least one month, ranges between 8.6 and 8.8. At the time of testing, the addition of 24 mg/L $Ca(OH)_2$ (the concentration tested here) raised the pH to 9.21.

To isolate the effect of the increased $Ca^{++}$ ion concentration by the addition of $Ca(OH)_2$, we calculated the concentration of $CaCl_2$ necessary to achieve an equimolar concentration of $Ca^{++}$ ions:

$$\frac{0.024\frac{g}{L}Ca(OH)_2}{74.093\frac{g}{mol}Ca(OH)_2} \times 110.985\frac{g}{mol}\ CaCl_2 = 0.036\frac{g}{L}\ CaCl_2$$

Accordingly, 36 g/L $CaCl_2$ was added to the dechlorinated tap-water used for exposure prior to testing on day 2 of testing in the $CaCl_2$ treatment group of focal females. This treatment does not alter the pH of the exposure water.

To isolate the effect of the pH elevation caused by the addition of $Ca(OH)_2$, we titrated to this pH using a saturated sodium hydroxide solution. This resulted in the addition of 15.4 mg/L NaOH to the dechlorinated tap-water used for exposure in the NaOH treatment group of focal females. Note this treatment does not affect $Ca^{++}$ ion concentration.

**Table S1. Responsiveness in all behavioral trials.** Response represents the number of responsive females over total females tested for each trial. P values are based on Fisher's exact test for count data.

| Exposure | Cue combination | Day | Treatment | Response | *p* |
|---|---|---|---|---|---|
| **$Ca(OH)_2$** | *X. birchmanni* vs *X. malinche* (conspecific vs heterospecific) | 0 | control | 7/14 | 0.1032 |
| | | | exposure | 12/14 | |
| | | 2 | control | 13/14 | 0.5892 |
| | | | exposure | 11/14 | |
| | | 4 | control | 13/14 | 1 |
| | | | exposure | 12/14 | |
| | | 12 | control | 10/14 | 0.1052 |
| | | | exposure | 14/14 | |
| **$CaCl_2$** | *X. birchmanni* vs *X. malinche* (conspecific vs heterospecific) | 0 | control | 14/16 | 1 |
| | | | exposure | 15/16 | |
| | | 2 | control | 14/16 | 1 |
| | | | exposure | 15/16 | |
| | | 4 | control | 14/16 | 1 |
| | | | exposure | 13/16 | |
| **NaOH** | *X. birchmanni* vs *X. malinche* (conspecific vs heterospecific) | 0 | control | 13/16 | 1 |
| | | | exposure | 14/16 | |
| | | 2 | control | 9/16 | 1 |
| | | | exposure | 9/16 | |
| | | 4 | control | 13/16 | 1 |
| | | | exposure | 12/16 | |
| **$Ca(OH)_2$** | *X. birchmanni* vs water (conspecific vs blank) | na | Control | 14/16 | 0.6539 |
| | | | exposure | 12/16 | |
| | *X. malinche* vs water (heterospecific vs blank) | na | control | 13/16 | 0.2258 |
| | | | exposure | 16/16 | |

**Table S2. Statistical analysis of net association time for each treatment and day for all trials in this study.** Comparison of mean net association time with olfactory cues. The test statistic, t, is reported for one-sample t-tests used when assumptions of normality were met. Where this was not the case Wilcoxon sign rank tests were substituted and the test statistic V is reported. df = degrees of freedom. Asterisks denote significant differences from zero (one-sample t-test, * $p < 0.05$, ** $p < 0.01$, *** $p < 0.001$).

| Exposure | Cue combination | Day | Treatment | df | test stat | *p* | |
|---|---|---|---|---|---|---|---|
| **$Ca(OH)_2$** | *X. birchmanni* vs *X. malinche* | 0 | control | 6 | t = 1.761 | 0.0644 | |
| | | | exposure | 12 | t = 2.104 | 0.0296 | * |
| | | 2 | control | 10 | t = 2.051 | 0.0314 | * |
| | | | exposure | 10 | t = -2.274 | 0.0463 | * |
| | | 4 | control | 12 | t = 2.708 | 0.0095 | ** |
| | | | exposure | 11 | t = -2.771 | 0.0182 | * |
| | | 12 | control | 9 | t = 1.898 | 0.0451 | * |
| | | | exposure | 13 | V = 94 | 0.0067 | ** |
| **$CaCl_2$** | *X. birchmanni* vs *X. malinche* | 0 | control | 13 | t = 6.251 | 1.5e-05 | *** |
| | | | exposure | 14 | t = 3.069 | 0.0042 | ** |
| | | 2 | control | 13 | t = 4.376 | 0.0004 | *** |
| | | | exposure | 14 | V = 53 | 0.7197 | |
| | | 4 | control | 13 | t = 3.928 | 0.0009 | *** |
| | | | exposure | 12 | t = 3.271 | 0.0067 | ** |
| **NaOH** | *X. birchmanni* vs *X. malinche* | 0 | control | 12 | t = 3.674 | 0.0016 | ** |
| | | | exposure | 13 | t = 4.517 | 0.0003 | *** |
| | | 2 | control | 8 | t = 2.457 | 0.0197 | * |
| | | | exposure | 9 | t = -3.284 | 0.0111 | * |
| | | 4 | control | 12 | t = 2.885 | 0.0068 | ** |
| | | | exposure | 11 | t = 3.993 | 0.0021 | * |
| **$Ca(OH)_2$** | *X. birchmanni* vs water | na | control | 13 | t = 3.2332 | 0.0065 | ** |
| | | | exposure | 11 | t = -0.9938 | 0.3417 | |
| | *X. malinche* vs water | na | control | 12 | t = -1.398 | 0.1874 | |
| | | | exposure | 15 | t = 2.979 | 0.0094 | ** |

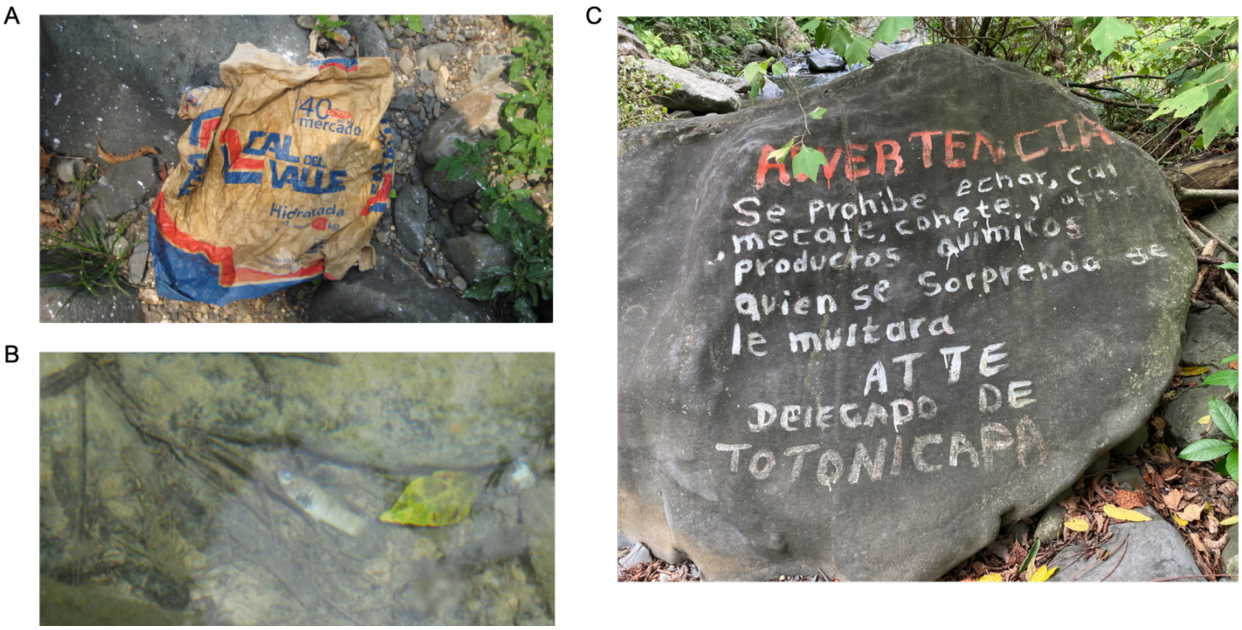


**Fig. S1.** Photographic evidence of calcium hydroxide usage in the Río Huazalingo, a natural stream containing *X. birchmanni* x *X. malinche* hybrids near the village of Totonicapa in the municipality of Tlanchinol in rural Hidalgo state, Mexico. A) An empty bag of calcium hydroxide found on the river bank near the site of an intentional introduction. B) A dead and decaying *Xiphophorus* found ~10 meters downstream of the empty bag in Fig. S2 A. C) Local signage prohibiting the use of lime (calcium hydroxide) among other things painted on a boulder next to the river. Text reads: "WARNING: It is prohibited to throw lime [$Ca(OH)_2$], *mecate* [plant-based toxin used for fishing], fireworks, and other chemicals. Whomever is caught will be fined. Sincerely, the [municipal] delegate of Totonicapa."

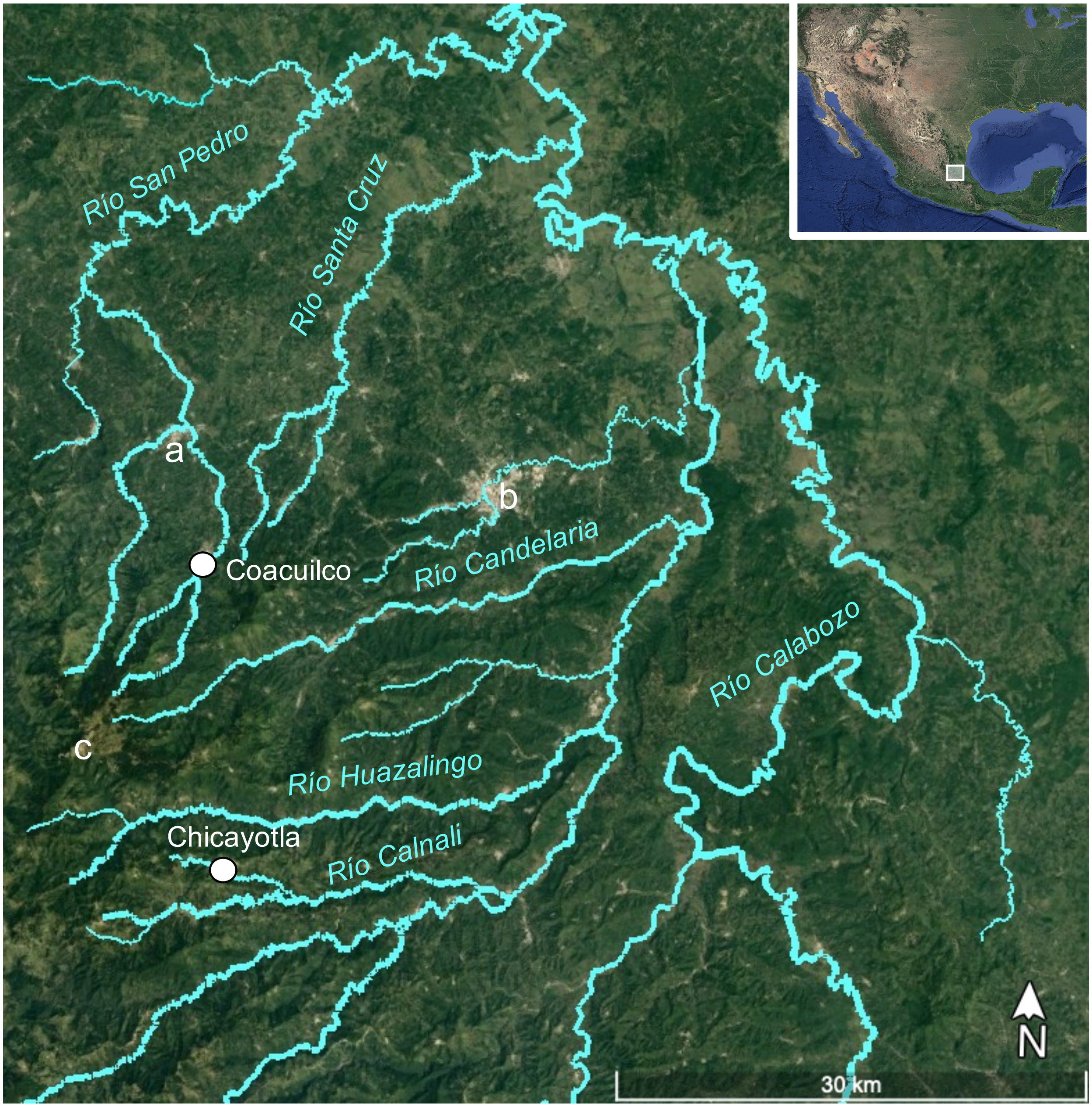


**Fig. S2.** Map of collection sites (white circles) for this study. Major river names are in turquoise. Major towns are marked by letters: a - San Felipe de Orizatlán, b - Huejutla de Reyes, c - Tlanchinol. Inset shows collection site location of *X. birchmanni* and *X. malinche* populations in Hidalgo, Mexico relative to a map of North and Central America. Image adapted from Google Earth.

**References**


1. McLennan DA, Ryan MJ. 1999 Interspecific recognition and discrimination based upon olfactory cues in northern swordtails. *Evolution* **53**, 880–888. (doi:10.2307/2640728)

2. McLennan DA, Ryan MJ. 1997 Responses to conspecific and heterospecific olfactory cues in the swordtail Xiphophorus cortezi. *Animal Behaviour* **54**, 1077–1088. (doi:10.1006/anbe.1997.0504)

3. Walling CA, Royle NJ, Lindström J, Metcalfe NB. 2010 Do female association preferences predict the likelihood of reproduction? *Behavioral Ecology Sociobiology* **64**, 541–548. (doi:10.1007/s00265-009-0869-4)

4. Fisher HS, Wong BBM, Rosenthal GG. 2006 Alteration of the chemical environment disrupts communication in a freshwater fish. *Proceedings of the Royal Society B: Biological Sciences* **273**, 1187–1193. (doi:10.1098/rspb.2005.3406)
